# *Mycoplasma gallisepticum* (MG) infection inhibits mitochondrial respiratory function in a wild songbird

**DOI:** 10.1101/2024.08.26.609819

**Authors:** Chidambaram Ramanathan, Elina Thomas, Amberleigh E. Henschen, James S. Adelman, Yufeng Zhang

## Abstract

An animal’s immune function is vital for survival, but some pathogens could manipulate their hosts’ immune and metabolic responses. One example is *Mycoplasma gallisepticum* (MG), which infects both the respiratory system and conjunctiva of the eye in house finches (*Haemorhous mexicanus*). MG has been shown to exhibit immune- and metabolic-suppressive properties, but the physiological mechanisms underlying these properties are still unknown. Recent studies demonstrated that mitochondria could serve as powerhouses for both ATP production and immunity, notably inflammatory processes, through regulating complex II and its metabolites. Consequently, in this study, we investigate the short-term (3d post inoculation) and long-term (34d post inoculation) effects of MG infection on the hepatic mitochondrial respiration of house finches from two populations and infected with two different MG isolates. After short-term infection, MG-infected birds had significantly lower state 2 and state 4 respiration, but only when using complex II substrates. After long-term infection, MG-infected birds exhibited lower state 3 respiration with both complex I and II substrates, resulting in lower respiratory control ratio compared to uninfected controls, which aligned with the hypothesized metabolic-suppressive properties of MG. Interestingly, mitochondrial respiration showed limited differences with house finch population of origin, MG isolate, and whether birds were recovered from infection or not. We propose that MG may target mitochondrial complex II for its immune-suppressive properties during the early stages of infection and inhibit mitochondrial respiration for its metabolic-suppressive properties at later stage of infection, both of which should delay recovery of the host and extend infectious periods.

## Introduction

When organisms face an infectious pathogen, they typically respond by activating innate immune defenses, including both rapid mechanisms and adaptive immune responses, which offers long-term, pathogen specific protection (1, 2). These immune activations and responses have been hypothesized to be costly in terms of both resource/nutritional allocation (3, 4) and metabolic demands (5, 6). Hosts could mitigate the cost of resource/nutrition by increasing food intake (7), however, the many metabolic costs from upregulating the immune system and production of inflammatory cytokines are likely unavoidable (8). Indeed, the energetic cost of immune response could range from upregulation of basal metabolic rates by 5%-15% after a mild pathogen infection (5, 9-12), to 25%-55% increases after severe infections (13-15). Because of the high metabolic demand required to kill or clear pathogens (known as resistance), organisms could adopt an alternative strategy to minimize harm from infection without necessarily reducing pathogen replication (known as disease tolerance) (16-19). Disease tolerance would allow hosts to limit the damage from infection, without bearing the large metabolic cost compared to resistance (20). In addition, pathogens also exhibit an extraordinarily wide range of mechanisms to manipulate host immunity by modulating host metabolism in order to facilitate and prolong infection and transmission (21), sometimes resulting in a decrease of metabolic rates (22). These different strategies employed by the host and the arms race between hosts and pathogens are fundamental parts of host-pathogen co-evolution (23).

Mitochondria have been demonstrated as a critical center of evolutionary tradeoffs between immunity and other life-history traits, including self-maintenance, as measured by immune defense (24-29). Mitochondria are best known as the primary sites of ATP production during aerobic respiration through oxidative phosphorylation (OXPHOS). Mounting an immune response could be energetically expensive, where the host mitochondria should become more efficient on ATP production following an immune challenge (30). Consistent with this idea, immune challenges typically increased mitochondrial respiration (30, 31), and treatment with a mitochondrial uncoupler reduced the pro-immune responsiveness against pathogens, such as skin swelling and developing fever (32, 33).

In addition to providing ATP to support immune function, mitochondria also tightly regulate a range of host immune responses, notably innate immune pathways, including RIG-I-like receptor (RLR) signaling, antibacterial immunity, and the sterile inflammatory response (27). Recent studies also documented that metabolites in the mitochondria, such as Krebs cycle intermediates, are vital signaling molecules for inflammatory response (34, 35). For example, succinate, a substrate for succinate dehydrogenase (SDH; mitochondria complex II), was shown to accumulate in macrophages during inflammation, acting as a signal to activate pro-inflammatory gene expression, notably IL-1β (35). Increased succinate oxidation by SDH is required to support the pro-inflammatory function of immune cells (36). Succinate is also released from cells and activates succinate receptor 1 (SUNCR1) to drive inflammatory pathways in other cells (37, 38). As infections and inflammation resolve, the pro-inflammatory effects of succinate could be counteracted by another metabolite, itaconate, which is derived from the Krebs cycle intermediate cis-aconitate (35). In sum, mitochondria provide not only energy, but also signals and reactions needed by the host immune response. Hence, mitochondria could be a central target for pathogen-driven manipulation of host immune responses (39-41).

Mycoplasma gallisepticum (MG) is bacterial pathogen that underwent a dramatic host shift from domestic poultry to wild birds, most notably house finches (*Haemorhous mexicanus*), during the early 1990s in the mid-Atlantic, USA. In house finches, MG causes severe conjunctivitis and decreases survival in the wild (42). Within the past three decades, MG first spread among house finches in eastern North America, eventually moving westward to the Pacific Northwest and Southwest of the USA (43, 44). Thus, MG endemism in house finch populations ranges from more than 20 years (such as Alabama) to 10-15 years (such as California), with MG was still not detected in Hawaiian Islands (16, 44). Moreover, since the host shift and the spread of MG in house finches are well-documented, this model serves as an ideal system to study host-pathogen interaction and coevolution across natural populations.

We have recently examined the role of mitochondrial respiratory function in this system, hypothesizing that MG infection should increase house finch mitochondrial respiration to cope with the elevated energy demand and to support pro-inflammatory responses. Surprisingly, we found that in Alabama, house finches captured during natural infection in the wild showed a trend toward lower state 3 respiration of liver mitochondria compared with birds that are MG-negative (45). These MG-positive house finches also showed significantly lower oxidative lipid and protein damage in liver tissue compared with the MG-negative counterparts (45). Although inconsistent with our original hypothesis of increased respiration, these observations are consistent with immune- and metabolic-suppressive properties of MG (45-47). Because the birds in that study were free-living, we could not assess their infection history, i.e., MG-positive animals could have been infected recently or infected for weeks (months is unlikely given MG-induced reduction in anti-predator responses (48)). Moreover, our previous study focused exclusively on Alabama house finches, which exhibit different responses to MG infection than do populations with a more recent history of MG endemism (16, 17, 23, 47, 49).

In this present study, we expand upon these results, asking whether MG-naïve individuals from two populations with different histories of MG endemism, Alabama (longer history) and California (shorter history), differ in both short- and long-term impacts of MG infection on mitochondrial respiration. Based on prior results, we hypothesized that MG-infected house finches would exhibit lower mitochondrial respiration compared to uninfected controls.

Moreover, we predicted that if MG-driven immunosuppression underlies these differences, California birds should show lower respiration during infection than Alabama birds, which have had a longer period of time to coevolve with the pathogen.

## Methods

### Ethics statement

All animal handling and euthanasia procedures were approved by the University of Memphis IACUC protocol (0842). All birds were captured and held under appropriate state and federal permits (AL: 2019107086868680, 2019107087668680, 2021122674268680, 2021122675668680; CA: S-190290001-19044-001; Tennessee Wildlife Resources Agency: 2252, 33080124, 35257117; United States Fish and Wildlife Service: MB82600B).

### Animal capture and housing

Juvenile house finches were captured by mist nets and feeder traps in Davis, California (CA) and Auburn, Alabama (AL) between July and August during 2019 and 2021, according to Henschen et al. (16). Briefly, only finches with no clinical signs of MG infection at the time of capture were transported back to the animal facility at the University of Memphis. All birds went through an acclimation and quarantine period after arrival (minimum of 40 days), including treatment with prophylactic medications to prevent disease from natural pathogens (16). Only individuals testing negative for anti-MG antibodies were included in the experiments (16). Finches were housed in medium flight cages (76 cm x 46 cm x 46 cm), and provided ad libitum water and food, consisting of a 20:80 mix of black oil sunflower seed:pellets (Roudybush Maintenance Nibles; Roudybush, Inc., Woodland, CA). We held light and dark cycles (12h:12h) and temperatures (∼22°C) constant.

### Experimental design

In the present study, we conducted both short (3-day infection; experiment 1, conducted in October, 2019) and long (34-day infection; experiment 2, conducted in October - November, 2019 and September - November, 2021) infection on house finches. In experiment 1, 10 birds from Alabama and 10 birds from California were randomly assigned into two groups, MG infected and control (Supplementary Table 1). We inoculated birds in MG infected group with 35 μL per eye of Frey’s medium containing a lower-virulence MG isolate (VA94 [7994–1 (6P) 9/17/2018] (50)) at high dose (7.5×10^6^ color changing units/ml (CCU/mL)). Control birds received sham-inoculations of 35 μL of Frey’s medium per eye. Three days post inoculation, all birds were euthanized as mentioned below.

Experiment 2 contained 44 birds from Alabama and 30 birds from California (Supplementary Table 1). Experimentally infected birds were inoculated with the same volumes as above using either a low dose (7.5×10^2^ CCU/mL) or high dose (7.5×10^6^ CCU/mL) of a lower-virulence MG isolate (VA94); or a low dose (7.5×10^2^ CCU/mL) of a higher-virulence isolate (VA13 [2013.089–15 (2P) 9/13/2013]). Control birds were mock inoculated with medium as above. Thirty-four days post inoculation (DPI), infected birds were further categorized into two groups: birds still exhibiting conjunctivitis (at 34 DPI) or pathogen load (at 28 DPI) were grouped as “chronic”, whereas birds without conjunctivitis (at 34 DPI) or pathogen load (at 28 DPI) were grouped as “recovered”. See “Statistical Analyses” regarding cases in which animals were positive for conjunctivitis, but negative for pathogen load and vice versa.

After each experiment, each bird’s conjunctiva was swabbed with a sterile cotton swab dipped in tryptose phosphate broth (TPB) and all samples were frozen for MG pathogen load quantification. Then birds were euthanized, and livers were quickly dissected out. The right lobe of the liver was placed in ice-cold mitochondrial isolation buffer (250 mM sucrose, 2 mM EDTA, 5 mM Tris-HCI, and 0.5% BSA, pH 7.4). The left lobe of the liver was snap-frozen in liquid nitrogen.

### Eye scores and pathogen load

Conjunctivitis in each eye was scored on a scale of 0–3 at 0.5 intervals according to Henschen et al. (16). Pathogen load was quantified via qPCR targeting the mgc2 gene according to Henschen et al. (16).

### Mitochondria isolation and respiration

Liver mitochondria were isolated from the right lobe of liver tissues according to Zhang et al. (45) and Hill et al. (51). Briefly, liver tissues were excised and minced in 10 mL isolation buffer (250 mM sucrose, 2 mM EDTA, 5 mM Tris-HCI, and 0.5% BSA, pH 7.4). Minced tissues were then homogenized in a Potter-Elvehjem PTFE pestle and glass tube. The resulting homogenate was centrifuged at 500 × *g* for 10 min at 4°C, and the supernatant filtered through cheesecloth before undergoing another round of centrifugation at 10,000 x g for 10 min. The resulting supernatant was discarded and the final mitochondria pellet was suspended in Mitochondrial Assay Solution (MAS-1: 2 mM HEPES, 10 mM KH2PO4, 1 mM EGTA, 70 mM sucrose, 220 mM mannitol, 5 mM MgCl2, 0.2% w/v fatty acid-free BSA, pH 7.4) according to Mookerjee et al. (52).

Liver mitochondria (0.35 mg/ml) respiration was measured in Mitochondrial Assay Solution (MAS-1) at 40°C using high resolution respirometry (Oroboros O2k, Innsbruck, Austria). For experiment 1, mitochondrial respiration was measured using both complex I (10 mM malate, 10 mM glutamate) and complex II (10 mM succinate, 2 μM rotenone) substrates in two chambers of the Oroboros concurrently. State 2 respiration was measured prior to addition of ADP. State 3ADP (maximal) respiration was induced by addition of 5 mM ADP, and state 4O (state 4Oligomycin; idling) respiration was induced by addition of 2 μg/mL oligomycin. Respiratory control ratio (RCR) was calculated by dividing state 3 ADP by state 4O respiration. For experiment 2, because of the large sample size, mitochondrial respiration when using complexes I and II substrates were sequentially measured in one chamber of the Oroboros. We measured mitochondrial respiration by adding complex I (10 mM malate, 10 mM glutamate) substrates (state 2) in the chamber. Then complex I substrates driven state 3ADP respiration was induced by the addition of 5 mM ADP and inhibited by addition of 2 μM rotenone. Complex II driven state 3ADP respiration was measured by addition of 10 mM succinate. In the end, state 4O respiration was measured by addition of 2 μg/mL oligomycin.

### Enzyme activity

Mitochondrial complex II activities were measured in the isolated liver mitochondria according to Spinazzi et al. (53). Complex II activities were measured in 25 mM potassium phosphate buffer contained 300 μM potassium cyanide, 20 mM succinate, 80 μM DCPIP (2,6-dichlorophenolindophenol), and 1mg/ml BSA at pH=7.5. Baseline activity was measured at 600 nm for 2 mins. The reaction was started by adding 50 μM decylubiquinone (DUB), mixed and measured for 3 min. The reaction was inhibited by adding 10 mM malonate. The citrate synthase (CS) activity was measured in the left lobe of liver homogenate and used as a proxy for mitochondrial density according to Spinazzi et al. (53). Briefly, 1 mg liver supernatant proteins were added to the buffer (100 mM Tris 0.1% (v/v) Triton X-100, 100 mM DTNB, and 300 mM acetyl-CoA at pH=8). Baseline activity was measured for 2 min, and reactions were started by adding the final concentration of 0.5 mM oxaloacetic acid and measured for 3 min at 412 nm. CS activity is widely used as a biomarker for mitochondrial content in tissues (54). Both absorptions were measured using Synergy H1 hybrid plate reader (Biotek, Winooski, VT, U.S.).

### Statistical analyses

All statistical tests were carried out using IBM SPSS, version 26.0. For experiment 1, we tested the effect of treatment and population (AL and CA) on mitochondrial respiratory markers and enzymatic activities using general linear models (GLMs) with these respiratory markers as dependent variables with treatment (control vs. MG) and population as main effects. For experiment 2, using GLMs, the year we conducted experiments had no significant effects on any respiratory markers (*P* > 0.08). In this experiment, however, conjunctivitis scores and pathogen loads would not always have resulted in the same categorization for a given bird as “recovered” or “chronic.” Specifically, 3 birds exhibited conjunctivitis on DPI 34, but no pathogen load quantified by qPCR on DPI 28, and 13 birds exhibited no conjunctivitis on DPI 34, but had positive pathogen load on DPI 38. If we categorized the birds using pathogen load alone, sample sizes for each group were as follows: control (18 birds), chronic (38 birds), and recovered (18 birds). If we categorized the birds using eye score alone, sample sizes for each group were as follows: control (18 birds), chronic (28 birds), and recovered (28 birds). As a result, we analyzed data using both grouping approaches. For both analyses, we tested mitochondrial respiration levels among birds that were chronic or recovered from MG with respiratory markers as dependent variables and population and infection status (control, chronic, or recovered) as main effects. Then, we tested different MG isolate and dosages using GLMs with respiratory markers as dependent variables, population and treatment (control, VA94 high dose, VA94 low dose, or VA13 low dose) as main effects. If a GLM showed significant effects of a given variable, Fisher’s LSD post hoc tests were performed. We considered *P* < 0.05 as statistically significant.

## Results

### *Experiment 1*: Short-term MG infection decreases mitochondrial complex II driven respiration

For short-term infection, population had significant effects on both state 2 (estimate (AL) = 1.75, *F*1,19 = 6.116, *P* = 0.026) and state 3ADP (estimate (AL) = 8.22, *F*1,19 = 17.046, *P* < 0.001) respiration with complex I substrates, in which Alabama birds exhibited higher state 2 and state 3ADP respiration than California birds (Supplementary Figure 1A). When comparing between MG infected vs non-infected birds, MG infected birds showed lower state 2 (estimate (MG infected) = -10.88, *F*1,19 = 4.735, *P* = 0.046) and state 4O (estimate (MG infected) = -17.61, *F*1,19 = 7.092, *P* = 0.018) respiration compared to their control counterparts only when using complex II substrates (Figure 1B). Low state 4O respiration resulted in higher complex II RCR (estimate (MG infected) = 1.12, *F*1,19 = 5.620, *P* = 0.032) for MG infected birds compared to the control (Figure 1C). The interaction between population and treatment was also significant for state 3 respiration using complex I substrates (estimate (AL × MG infected) = -20.01, *F*1,19 = 5.596, *P* = 0.032), such that Alabama control birds showed higher respiration than Alabama MG infected, California control and MG infected. We detected no major differences in the enzyme activity of citrate synthase (CS) and complex II between the populations or treatment groups (Figure 1D, *P* > 0.553). These results indicated that short-term infection decreases complex II driven respiration without affecting mitochondria density.

**Figure 1.**
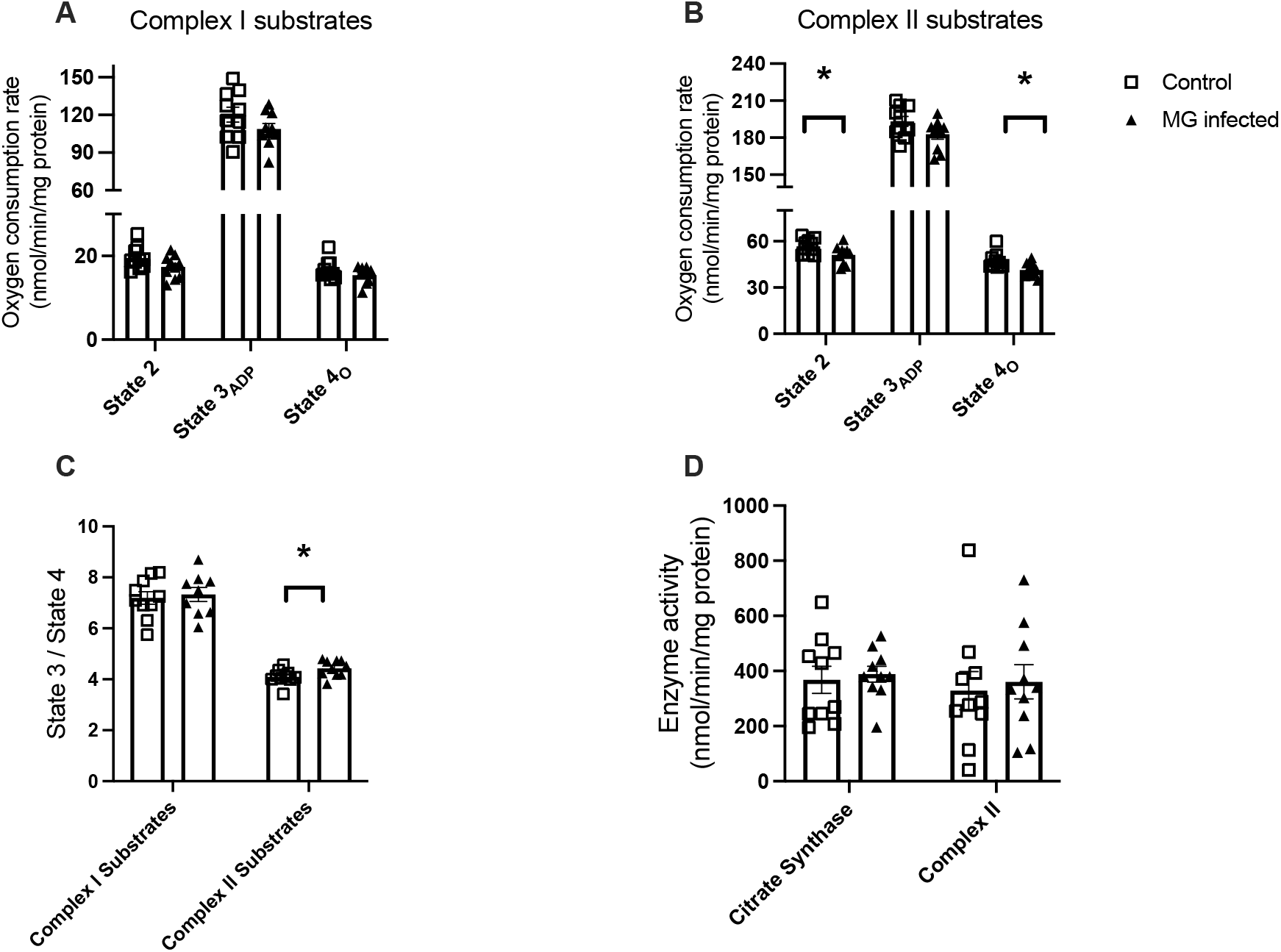
Short-term infection of MG on house finch reduced mitochondrial respiration driven proton leak using complex II substrates. Mitochondrial respiration and enzymatic activity from isolated liver mitochondria for birds three-day post inoculation with *Mycoplasma gallisepticum* (MG infected) and uninoculated birds (Control). Bar graphs showing the states 2, 3, and 4 respiration rates with complex I (10 mM malate, and 10 mM glutamate (A)), complex II (10 mM succinate with 2 μM rotenone (B)), respiration control ratio (RCR (C)), and citrate synthase activity in liver homogenate and complex II activity in isolated liver mitochondria (D). Squares represent uninfected birds, and triangles represent MG-infected birds. Data are presented as mean ± SEM. * indicates P < 0.05

### *Experiment 2*: Long-term MG infection decreases mitochondrial complex I and II driven respiration

For long-term MG infection (34 days post inoculation), categorizing birds into chronic and recovered by either pathogen load or eye score showed very similar results. Results based on eye score were included in the Supplementary Figure 2. When we group birds by pathogen load, population did not affect any mitochondrial respiratory markers (all *P* > 0.292). However, control birds had higher state 2 (estimate (control) = 8.73, *F*2,62 = 36.456, *P* < 0.001) and state 3ADP for both complex I (estimate (control) = 24.53, *F*2,62 = 26.224, *P* < 0.001) and complex II driven respiration compared to either chronic or recovered birds (estimate (control) = 91.82, *F*2,62 = 93.144, *P* < 0.001), but state 4O respiration was not significantly affected (*P* = 0.151, Figure 2A and B). Consequently, RCR for both complex I (estimate (control) = 0.38, *F*2,62 = 6.305, *P* = 0.003) and complex II driven (estimate (control) = 1.61, *F*2,62 = 27.330, *P* < 0.001) respiration were also higher for control birds than either chronic or recovered counterparts (Figure 2C).

**Figure 2.**
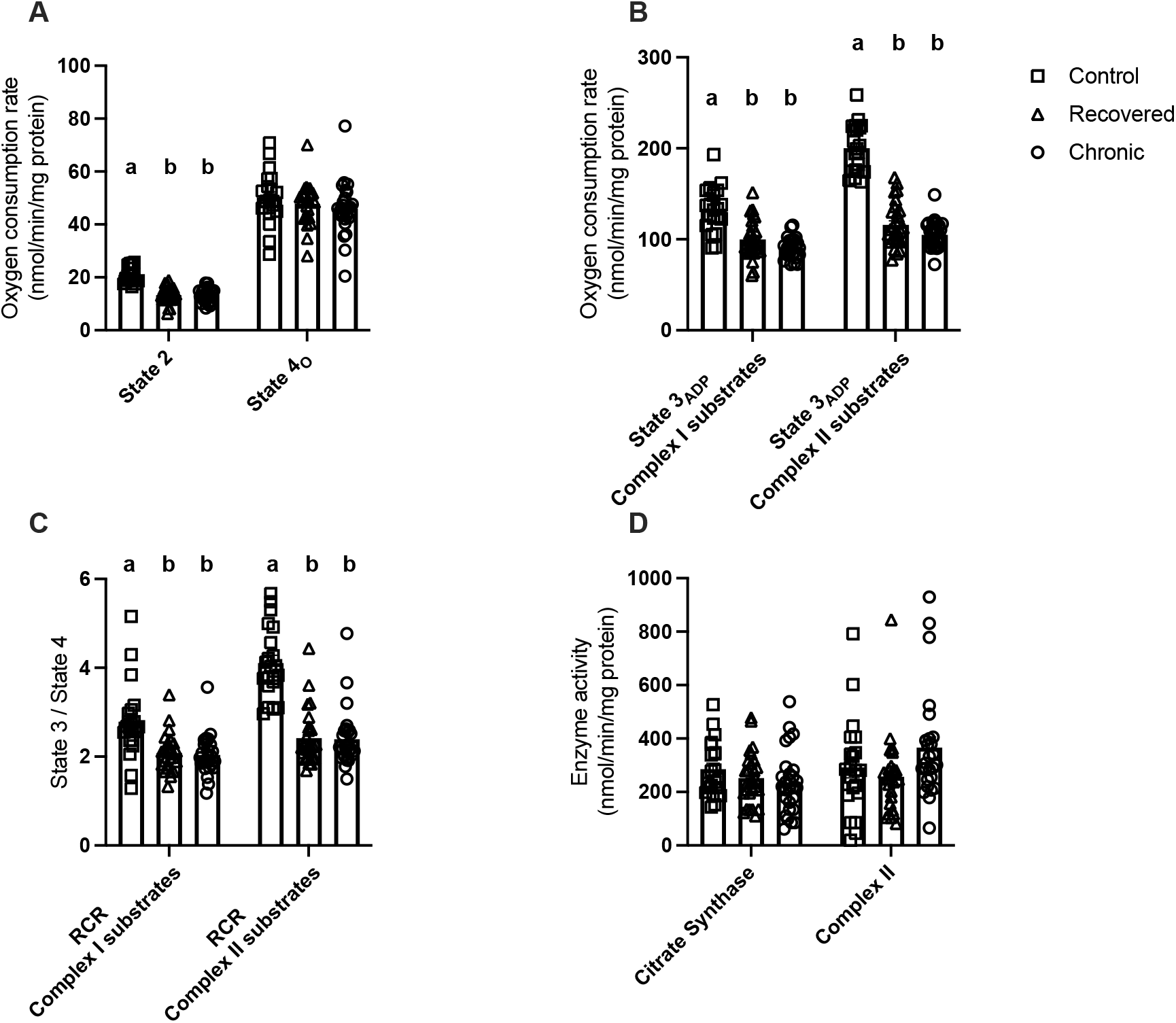
Long-term infection of MG on house finch reduced mitochondrial complexes I and II respiration. Mitochondrial respiration and enzymatic activity from isolated liver mitochondria for birds twenty-eight days post inoculation with recovered (no pathogen load), chronic (positive pathogen load), and uninoculated birds (Control). Bar graphs showing that states 2 and 4O respiration rates (A) and state 3ADP respiration with complex I (10 mM malate, and 10 mM glutamate) and complex II (10 mM succinate with 2 μM rotenone (B) substrates; respiration control ratio (RCR) (C) citrate synthase activity in liver homogenate and complex II activity in isolated liver mitochondria (D). Squares represent uninfected birds, triangles represent recovered birds, and circles represent chronic birds. Data are presented as mean ± SEM. Bars that do not share a letter depict means that are significantly different from one another (P < 0.05).

When comparing between different MG isolates at 34 DPI, state 3ADP for both complex I (estimate (control) = 35.31, *F*2,62 = 15.886, *P* < 0.001) and II driven (estimate (control) = 97.34, *F*2,62 = 66.101, *P* < 0.001) respiration were significantly lower for MG infected birds compared to control regardless of MG isolates (Figure 3B). This decrease in state 3ADP respirations also resulted in lower RCR for both complex I (estimate (control) = 0.58, *F*2,62 = 4.477, *P* = 0.007) and II driven (estimate (control) = 1.71, *F*2,62 = 18.401, *P* < 0.001) respiration, with stable state 4O respiration (*P* = 0.334) (Figure 3 C). State 2 respiration (estimate (control) = 9.69, *F*2,62 = 28.931, *P* < 0.001) also differed between groups (Figure 3 A). Specifically, control birds had higher state 2 respiration compared to low-dose/high-virulence (VA13; *P* < 0.001), and low-dose/low-virulence (VA94; *P* < 0.001) and high-dose/low-virulence (VA94) (*P* < 0.001). Both low-dose/high-virulence (VA13; *P* = 0.015) and high-dose/low-virulence (VA94; *P* = 0.041) infection also resulted in higher state 2 respiration compared to low-dose/low-virulence (VA94). The interaction between population and treatment interaction also helped explain variation in complex II driven state 3ADP respiration (*F*2,62 = 2.845, *P* = 0.046), with Alabama birds have higher state 3 respiration than California birds when infected with low-dose/low-virulence (VA94) MG (*P* = 0.023) (Data not shown). These results indicated that long-term MG infection decreases both mitochondrial respirations regardless of MG isolates and the status of the infection.

**Figure 3.**
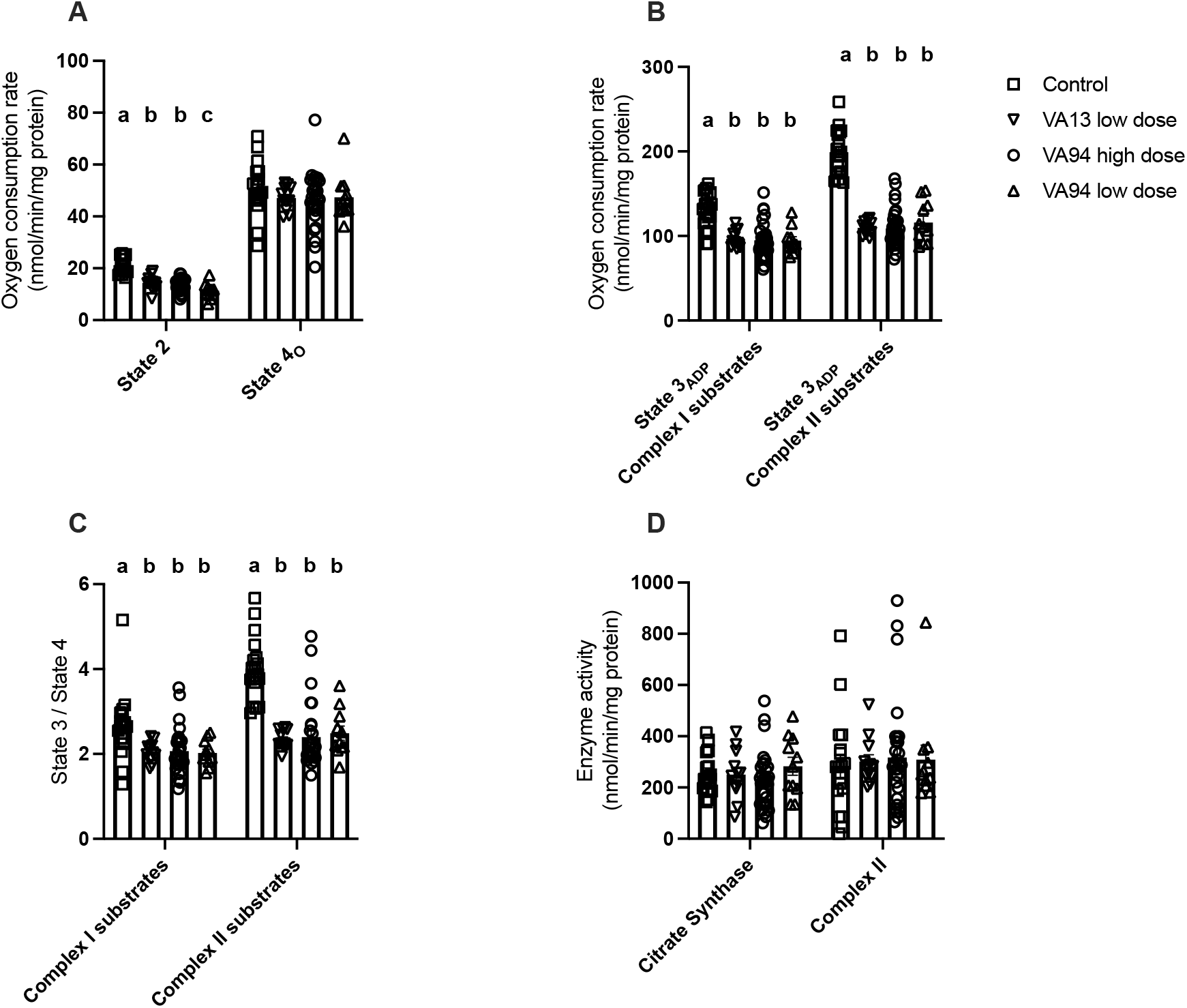
Long-term infection of MG on house finch reduces mitochondria respiration regardless of the virulence of the MG isolates. Mitochondrial respiration and enzymatic activity from isolated liver mitochondria from birds twenty-eight days post inoculation with VA13 low dose (7.5×10^2^ CCU/mL), VA94 high dose (7.5×10^6^ CCU/mL), VA94 low dose (7.5×10^2^ CCU/mL) and uninoculated birds (Control). Data include states 2 and 4O respiration rates; A) and state 3ADP respiration with complex I (10 mM malate, and 10 mM glutamate) and complex II (10 mM succinate with 2 μM rotenone; B) substrates; respiration control ratio (RCR; C); citrate synthase activity in liver homogenate and complex II activity in isolated liver mitochondria (D). Squares represent uninfected birds, down-pointing triangles represent VA13 low dose birds, and circles represent VA94 high dose birds, up-pointing triangles represent VA94 low dose birds. Data are presented as mean ± SEM. Bars that do not share a letter depict means that are significantly different from one another (P < 0.05).

## Discussion

The present study investigated the short- and long-term effects of MG infection on mitochondrial respiratory function in the liver of house finches. The liver acts as the metabolic hub in organisms and plays a central role in both metabolism and immune defense against invading pathogens (55). Short-term MG infection in house finches showed reduced complex II states 2 and 4 and respiration, but not complex I mediated respiration. On the other hand, long-term infection reduced levels of both complex I and complex II driven respiration compared with control birds. Interestingly, birds that were recovered from MG infection with no symptoms and pathogen load (recovered) showed mitochondrial respiration similar to birds that were still exhibiting symptoms (chronic). However, complex II enzymatic activities were not changed for either short- and long-term infection, even when complex II driven respiration was reduced. Mitochondrial densities, represented by citrate synthase activity, also remained unchanged during infection. These observations suggested that MG might exhibit metabolic suppressive properties, via interference with mitochondrial function.

For short-term infections, measurements were conducted 3-days post inoculation (DPI). On DPI 3, infected house finches started to show eye lesions (17), and infected birds can excrete the bacteria and become infectious starting at DPI 3 (56). More importantly, cDNA microarrays data suggested that genes associated with immunity, metabolism, and redox status were differentially expressed in spleens between infected finches compared to their controls (46). These observations during the early stage of MG infection suggested that early acting innate immune processes have been activated at DPI 3, and the metabolic and immune profile of the organisms remain relatively similar for 2 weeks after infection. For our birds, liver mitochondrial state 2 and state 4o respiration levels were reduced on DPI 3 for infected birds compared to their controls only when using complex II substrates. State 2 and state 4o respiration, are measured under conditions where mitochondria are provided no external ADP for production of ATP (state 2) or ATP synthase are inhibited (state 4o), so oxygen consumption comes from proton leakage across the inner mitochondrial membrane. Even though state 4o respiration changes disproportionately with changes of proton motive force (pmf), but these changes of state 4o respiration could be explained by proton leak across the mitochondrial inner membrane (57), making state 4o respiration as an indirect measurement of proton leak rate (58). Consequently, this suggested that MG infected birds have lower mitochondria proton leak rate compared to the controls only when using complex II substrates on DPI 3. Reduced state 4o respiration resulted in the higher complex II driven RCR for MG infected birds, which indicated that the short-term infected birds’ liver mitochondria are more coupled during early infection.

It is very interesting that the differences in mitochondria proton leak were only observed when using complex II substrates, not complex I substrates, between MG infected and controls on DPI 3. It is unlikely that complex I and complex II substrates affected the mitochondrial inner membrane proton permeability differently, whereas the fundamental differences between complex I and complex II driven respirations might offer some insights. Succinate oxidation (complex II) would generate higher pmf than NADH-linked (complex I) substrates, mostly due to the differences in redox potentials and thermodynamic “gearing” (59). Consequently, biological regulation of succinate oxidation *in vivo* is extensive and highly tunable, much more than NADH-linked substrates (34, 60). This tight control of succinate oxidation might be required to maintain mitochondria pmf within a range that allows optimal ATP production and signaling (by ATP/ADP, O2•−/H2O2, metabolite levels) but minimizes oxidative damage by O2•−/H2O2 (59). Recently, succinate oxidation has been shown to serve as a vital regulator in variety of biological processes ranging from ischemia (61), tumorigenesis (62), epigenetic control (63), to inflammation (36). More specifically, on immunity, increased succinate oxidation with elevated mitochondrial membrane potential drives pro-inflammatory responses (such as IL-1β secretion) in macrophages, whereas inhibition of succinate oxidation would promote anti-inflammatory responses (i.e. IL-10 secretion (24, 36)). As a result, SDH could be a prime target for pathogens to influence the immune response of the hosts.

*Mycoplasma* bacteria are known for manipulating host immune defense system (64). Similarly, MG infection could cause the suppression of certain immune components, such as down-regulation of pro-inflammatory cytokines (IL-1β, IL-8, CCL20, and IL-12), especially at the early stage of infection in chicken (65). In house finch, certain pro-inflammatory cytokines, such as IL-6 and IL-1β, were upregulated in conjunctiva and eyelid after MG infection (66). On the other hand, IL-10, which serve as an anti-inflammatory cytokine, was also upregulated during early MG infection (17, 66). But these changes can depend on the house finch population, MG isolates, and infection dosages. Higher-virulence, higher dosages, and evolutionarily derived isolates tended to induce higher levels of pro-inflammatory cytokines compared to isolates which are original, lower-virulence, and lower dosages (67). House finches from different populations also exhibited different inflammation profiles after MG infection. Eastern birds, who has been co-evolved with MG longer than western birds, exhibited low/no changes of pro-inflammatory cytokines (such as IL-1β) with upregulation of anti-inflammatory cytokines (such as IL-10) levels at early stages of infection (17, 46, 66, 67). For the present study, we used lower-virulence isolates with relatively high dosages (VA94 at 7.5×10^6^ CCU/mL) during short-term infection. Previous studies using the same isolates and dosages indicated lower pro-inflammatory responses with upregulation of anti-inflammatory cytokines in eastern birds compared to their western counterparts (16). Even though Alabama birds showed higher complex I driven state 2 and 3 respirations in this study, these differences were largely due to control birds, whereas mitochondrial respiration was relatively similar between these two populations after MG infection (*P* > 0.282). Taken together, these results suggest that MG may have immunosuppressive properties in house finches, and such immunosuppression could be accomplished by targeting the SDH (complex II) of the host.

The potential mechanisms of MG targeting SDH are currently unknown. However, SDH enzymatic activities in this study were not affected by MG infection even with lower mitochondrial respiration using complex II substrate (Figure 1D). It is worth noting that SDH enzymatic activity assay measures the maximal activities of SDH, and that maximal respiration (state 3) of complex II substrates were also similar between infected birds and controls. This suggests the inhibition of complex II driven respiration likely happened on the metabolite level of the host cells. It has been shown that when another strain of Mycoplasma, *Mycoplasma arginine*, infects VM-M3 cancer cells, succinate production from cell mitochondria increases and succinate is eventually exported into the extracellular medium (68). This increase of succinate production and release correlates to increases in other metabolites, including intracellular level of itaconate (68). Itaconate, then, can further inhibit SDH activity which serves as an anti-inflammatory signal of the host (35). Specifically for MG, metabolomics have indicated that TCA cycle metabolites, such as fumarate and succinate, were detected in MG and other strains of mycoplasma after infection (69, 70). This is extremely surprising because none of these mycoplasma genomes contain enzymes for a TCA cycle, suggesting that these metabolites have been up-taken from the medium (70). More studies are needed to unveil the interesting and vital mechanisms of the immunosuppressive properties of MG.

For long-term infection, we conducted measurements on DPI 34. At this stage, 16 Alabama birds exhibited no pathogen load (recovered), whereas 22 Alabama birds still contain pathogen (chronic), based on pathogen detection via qPCR. For California birds, only 2 were recovered, but 16 birds still have pathogen load. Surprisingly, mitochondrial respiration levels were not different between recovered or chronic groups, though infection decreased state 3 respiration for both complex I and II substrates. This is consistent with the previously documented metabolic suppressive properties of MG. For example, microarray data showed decreased expression of genes regulating metabolism, such as prosaposin and spermidine/spermine N1-acetyltransferase, two weeks after infection (49). The cDNA macroarray also indicated that a number of genes associated with redox status and the electron transport system, such as cytochrome oxidase subunit I and III and NADH dehydrogenase subunit 4, are downregulated in the MG-infected house finches (71). It is important to note here that MG increases body temperature and resting metabolic rates in house finches early in infection, but these approach control levels after 14 days (17, 72). Similarly, we only observed lower maximal mitochondrial respiration after long-term infection in the present study, suggesting MG targeting of immune responses early in infection, and the host’s metabolic processes later in infection (16).

Surprisingly, we did not observe any differences in population and MG isolates on mitochondrial respiration after infection. Prior works suggested that western birds exhibit more pronounced immune- and metabolic-suppression than their eastern counterparts, in terms of systemic, i.e. splenic, gene expression (46, 49, 67). California house finches, who have coevolved with MG for 10-20 years; and Alabama birds, coevolved for 20-25 years were employed for this study (16). It is possible that these temporal differences might not be large enough to observe any differences on mitochondrial respiration levels in the liver. The dosages and virulence of MG isolates also exhibited limited differences after infection in present study, even though high dose high virulence isolates are more likely to cause infection and related symptoms (73). Still, and regardless of population and MG isolates, long-term MG infection decreased the mitochondrial respiration, even among birds recovered from MG. This is particularly important because it suggests that MG infection could have long-term organismal, and potentially ecological, consequences even after birds recover. MG infection increases the mortality rate of house finches in the wild (42, 74), not necessarily through directly killing the host (50, 75, 76), but more likely by increasing the risk of mortality through indirect mechanisms, such as reduced locomotion and anti-predator behaviors (48). Especially since these reductions in anti-predator defense can occur during infection but in the absence of conjunctivitis (48), it is possible that they result from the metabolic effects of MG (48). Consequently, birds could still exhibit the same behavioral effects, which lead to higher mortality, even after recovery from MG (48).

In conclusion, we have shown that MG infection suppressed the hepatic mitochondrial respiration in house finches. These changes are consistent with previous studies, and we propose that the immune- and metabolic-suppressive properties of MG may be accomplished by targeting mitochondria of the host. During early infection, MG targets the SDH of the host to suppress its pro-inflammatory responses (36). Furthermore, MG decreases mitochondria respiration and coupling efficiencies, which suppress the metabolism of the host later during infection. Unfortunately, the exact underlying mechanisms for such suppressions remain unknown. Future mechanistic studies using *in vitro* models could shed light on the proposed pathway. Moreover, a detailed longitudinal monitoring of the metabolic and immune responses of the host after MG infection is also needed. Together, integrating *in vitro, ex vivo*, and *in vivo* approaches could help us better understand how such physiological mechanisms impact the arm race between hosts and pathogens.

## Acknowledgement

We would like to thank Matthew Butawan, Francis Edward Tillman Jr, and Thomas Lackie for their technical support. This research was funded by National Science Foundation grants IOS2224556 and IOS2037735 to Y.Z. and IOS1950307 to J.S.A.

## Author Contributions

C.R., A.E.H, J.S.A., and Y.Z. contributed to conceptualization, methodology, investigation, data curation, visualization, supervision, writing and editing. E.T. contributed to investigation and editing.

