## Supplementary material for "*Mycoplasma gallisepticum* (MG) infection inhibits mitochondrial respiratory function in a wild songbird": Supplmental figures

Supplementary table 1. Sample size for each experiment: chronic (positive pathogen load), recovered (no pathogen load), AL: Alabama, CA: California.

| Experiment | Groups | Population | Sample size |
| --- | --- | --- | --- |
| 3-day post inoculation | Control | AL | 5 |
|  |  | CA | 5 |
|  | VA94 high | AL | 5 |
|  |  | CA | 5 |
| 34-day post inoculation | Control | AL | 6 |
|  |  | CA | 12 |
|  | VA94 high chronic | AL | 17 |
|  |  | CA | 7 |
|  | VA94 high recovered | AL | 9 |
|  |  | CA | 0 |
|  | VA 94 low chronic | AL | 0 |
|  |  | CA | 4 |
|  | VA 94 low recovered | AL | 6 |
|  |  | CA | 1 |
|  | VA13 low chronic | AL | 5 |
|  |  | CA | 5 |
|  | VA13 low recovered | AL | 1 |
|  |  | CA | 1 |

Supplementary Figure 1. Mitochondrial respiration and enzymatic activity from isolated liver mitochondria for house finches three-day post inoculation between Alabama and California birds. Bar graphs showing the states 2, 3, and 4 respiration rates with complex I (10 mM malate, and 10 mM glutamate (A)), complex II (10 mM succinate with 2 μM rotenone (B)), respiration control ratio (RCR (C)), and citrate synthase activity in liver homogenate and complex II activity in isolated liver mitochondria (D). Square represent Alabama birds, and triangles represent MG-infected birds. Data are presented as mean ± SEM. * indicates P < 0.05

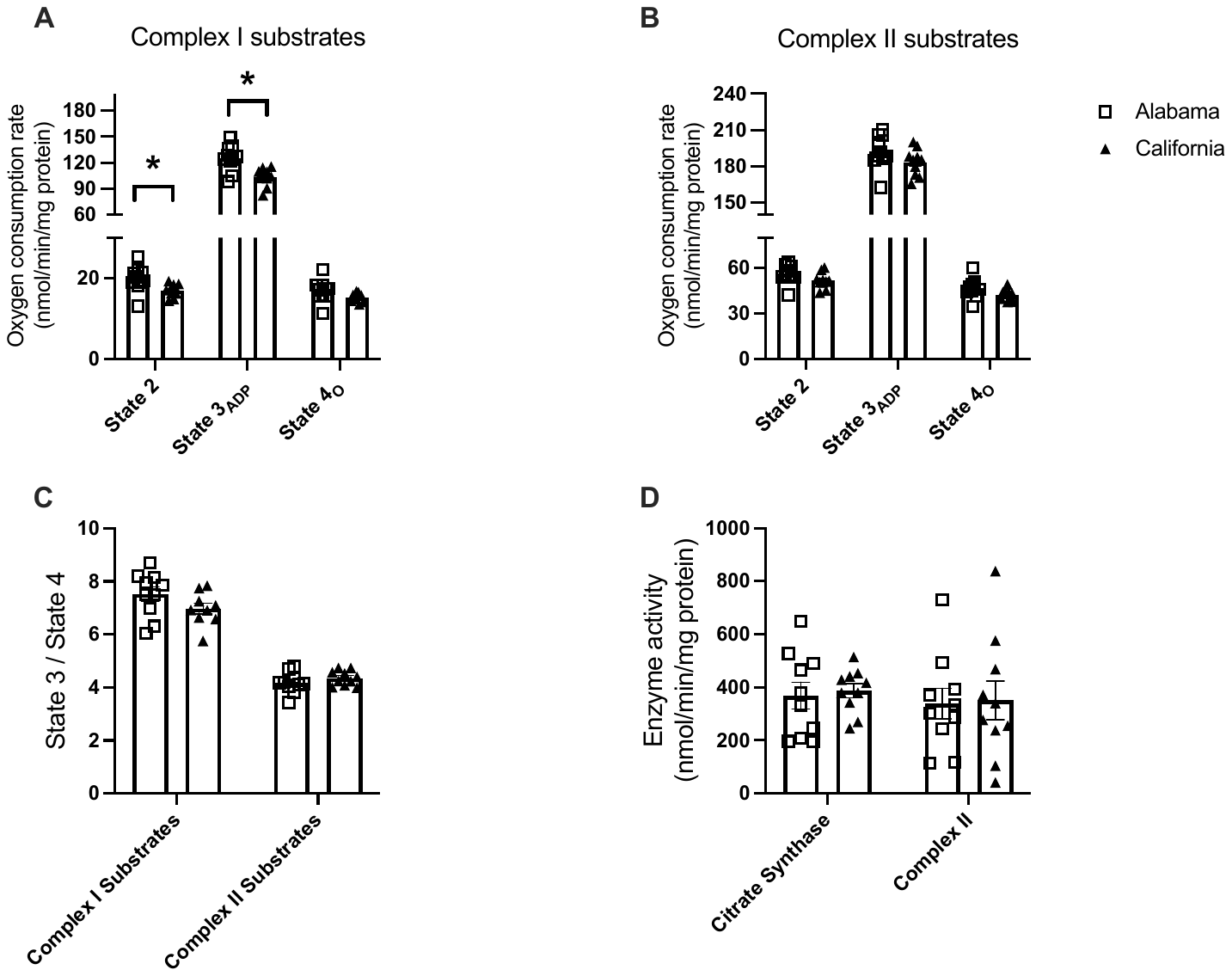

Supplementary Figure 2. Mitochondrial respiration and enzymatic activity from isolated liver mitochondria for birds twenty-eight days post inoculation with recovered (no symptom), chronic (still exhibit symptom), and uninoculated birds (Control). Bar graphs showing that states 2 and 4_O_ respiration rates (A) and state 3_ADP_ respiration with complex I (10 mM malate, and 10 mM glutamate) and complex II (10 mM succinate with 2 μM rotenone (B) substrates; respiration control ratio (RCR) (C), citrate synthase activity in liver homogenate and complex II activity in isolated liver mitochondria (D). Squares represent uninfected birds, triangles represent recovered birds, and circles represent chronic birds. Data are presented as mean ± SEM. Bars that do not share a letter depict means that are significantly different from one another (P < 0.05).

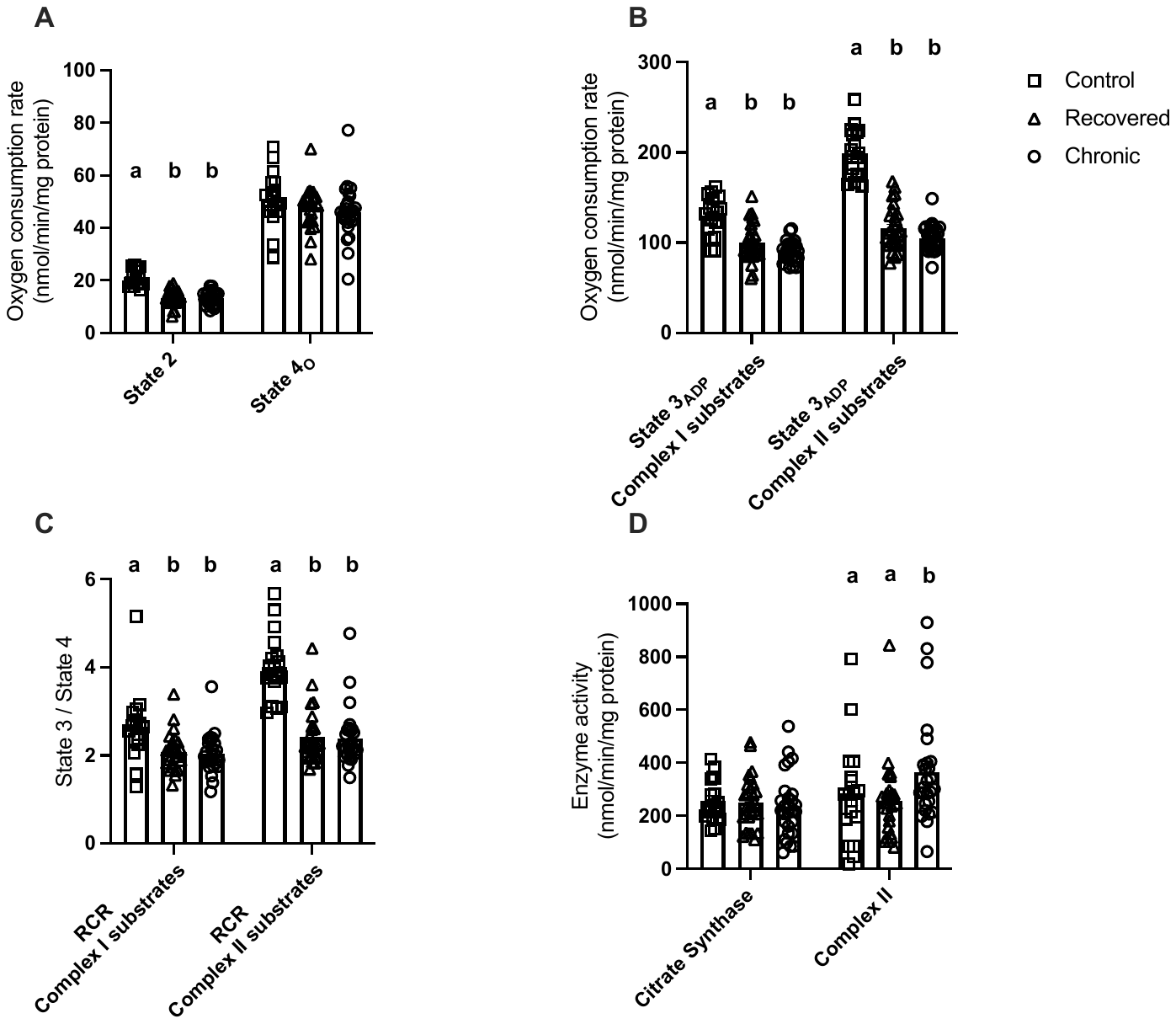
